# Cross Potential Selection: A Proposal for Optimizing Crossing Combinations in Recurrent Selection Based on the Ability of Future Inbred Lines

**DOI:** 10.1101/2024.04.05.588296

**Authors:** Kengo Sakurai, Kosuke Hamazaki, Minoru Inamori, Akito Kaga, Hiroyoshi Iwata

**Author notes:** Corresponding author: Hiroyoshi Iwata.

## Abstract

In plant breeding programs, rapid production of novel varieties is highly desirable. Genomic selection allows the selection of superior individuals based on genomic estimated breeding values. However, it is worth noting that superior individuals may not always be superior parents. The choice of the crossing pair significantly influences the genotypic value of the resulting progeny. This study introduced a new strategy for selecting crossing pairs, termed Cross Potential Selection (CPS), designed to expedite the production of novel varieties. The CPS assesses the potential of each crossing pair to generate a novel variety. It considers the segregation of each crossing pair and computes the expected genotypic values of the top-performing individuals, assuming that the progeny distribution of genotypic values follows a normal distribution. We simulated a 10-year breeding program to compare CPS with three other selection strategies. CPS consistently demonstrated the highest genetic improvements among the four strategies in early cycles. In particular, during the middle cycles of the breeding program, CPS exhibited the highest genetic improvement of 73% of the 300 independent breeding simulations. In a long-term breeding scheme, some progeny distributions of genotypic values may deviate from normal distribution, affecting the efficiency of CPS. Nevertheless, compared with the other three strategies, CPS achieved significant short-term genetic improvements. In conclusion, CPS holds substantial promise for enhancing the efficiency of plant breeding programs.

**Article Summary:** This study introduces a novel plant breeding strategy termed Cross Potential Selection (CPS), which was designed to expedite the production of novel varieties. The CPS evaluates the potential of each crossing pair for the target generation. Through comparative breeding simulations, CPS demonstrated superior performance over the three alternative breeding strategies, particularly in the early cycles. These findings suggest that CPS holds significant promise for enhancing plant breeding efficiency.

## INTRODUCTION

Plant breeding aims to enhance the genotypic value of a target trait through selection and crossing. The process of selecting candidates for plant breeding is important, because it directly influences the outcomes of breeding programs. Genomic prediction (GP) models have been developed to aid in selecting superior candidates using genome-wide polymorphism data (Meuwissen et al., 2001). These models were constructed using training data, comprising genome-wide marker and phenotypic data, to estimate the effects of markers across the genome on a target trait. By leveraging GP models, the genomic estimated breeding values (GEBVs) can be obtained for novel genotypes without the need to conduct field trials. This approach, known as genomic selection (GS), enables the selection of superior candidates based on the GEBVs.

In plant breeding, once candidates are selected, the next step involves determining the crossing pairs to generate progeny. As crossing pairs directly influence the genotypic values of the progeny, various crossing strategies have been devised to select crossing pairs from the current population. Genetic diversity plays a crucial role in driving genetic improvements in breeding programs (Sanchez et al., 2023, 2024). Some crossing strategies consider factors, such as the expected genotypic values of the next generation and the degree of kinship among the selected individuals (Akdemir & Sánchez, 2016; Kinghorn, 2011). In addition, the usefulness criterion (UC) was introduced as a selection index for crossing pairs (Schnell & Utz, 1975). UC represents the expected value of promising individuals when producing recombinant inbred lines (RILs) from each crossing pair. It is defined as *UC* = *μ* + *ihσ*, where *μ* is the mean genotypic values of the cross, *i* is the selection intensity, *h* is the square root of heritability, and *σ* is the square root of genetic variance for the RILs. In a two-way cross, the genetic variance of inbred progenies can be computed using the recombination rates between two markers, estimated marker effects for the target trait, and the marker data of each cross (Lehermeier et al., 2017). Allier et al. (2019) extended this formula to three- and four-way crosses. In breeding simulations, genetic improvements in the F_5_ population produced from crosses selected using UC were greater than those produced from crosses selected using GEBVs (des Déserts et al., 2023). Crossing selection based on UC has significant potential for enhancing genetic improvement in plant breeding programs.

Gaynor et al. (2017) introduced a two-part strategy to enhance the efficiency of plant breeding programs. This strategy comprises a “product development component,” which identifies promising individuals for release as varieties, and a “population improvement component,” which boosts the genotypic values of the breeding population through rapid recurrent genomic selection conducted twice a year. Optimal cross selection (OCS) employs the crossing strategy devised by Kinghorn (2011) for population improvement component in a two-part strategy (Gorjanc et al., 2018). OCS has demonstrated substantial long-term genetic improvements compared to the standard two-part strategy that selects candidates based on GEBVs. However, only a few breeding strategies have surpassed GS in achieving higher short-term genetic improvements. Allier et al. (2019) employed UC in their breeding scheme and achieved higher short-term genetic improvement than GS. However, their breeding scheme was limited to crop species capable of producing doubled haploids. Furthermore, rapid recurrent selection was not employed in their breeding scheme, as their focus was not on expediting variety development. In practical breeding scenarios, it is necessary to expedite production of novel varieties. Therefore, it is imperative to develop breeding strategies capable of achieving significant short-term genetic improvements to rapidly produce novel varieties. However, such breeding strategies are yet to be developed. This study introduced a novel breeding strategy for the rapid production of novel varieties. We proposed a new strategy termed Cross Potential Selection (CPS), which integrates the UC and a two-part strategy. To assess the efficacy of CPS, we conducted 300 independent breeding simulations based on a two-part strategy, comparing the genetic improvements of the four breeding strategies: GS, OCS1, OCS2, and CPS. Additionally, we analyzed the genetic variance and fixation of beneficial alleles within the population improvement component across strategies to delineate the unique characteristics of CPS. To focus on the differences in improvement efficiency among the selection strategies, we conducted comparisons assuming that the marker effects were known rather than estimated.

## MATERIALS AND METHODS

### Population in breeding simulations

The population used in breeding simulations was generated using whole-genome sequence data from a diverse panel of 198 soybean accessions. These accessions primarily comprise the Japanese and global soybean mini-core collections (Kaga et al., 2011; Kajiya-Kanegae et al., 2021). The whole-genome sequence data encompassed 4,776,813 single-nucleotide polymorphisms (SNPs) distributed across 20 pairs of chromosomes. SNPs that were heterozygous or had >95% missing data were excluded, along with those with a minor allele frequency <0.1. Additionally, SNPs were filtered based on linkage disequilibrium, with pairs <0.6 selected, resulting in a final set of 61,426 SNP markers. From a pool of 61,426 SNPs, we randomly selected 4,000 SNPs (200 SNPs per chromosome) for each independent breeding simulation to conserve computer memory and accelerate the simulations. We assumed that each chromosome had a length of 1 Morgan and that there was a linear relationship between map distances and physical distances. Consequently, the linkage map positions were calculated based on the physical positions of adjacent SNPs. The simulation setup was adopted from Diot & Iwata (2023).

In this breeding simulation, of the 4,000 SNPs, 1,000 were randomly designated as quantitative trait nucleotides (QTNs) and non-zero effects were assigned to the genotypes of the initial population. The effects of QTNs followed a multivariate normal distribution.

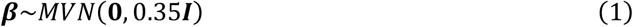

where ***β*** represents the vector of the effect for the 1,000 QTNs, and ***I*** denotes the identity matrix. The effect of the remaining 3,000 SNPs were set to zero.

Each breeding simulation commenced with 150 individuals, generated from a four-way cross. A diverse panel of 198 soybean accessions was divided into four groups based on 4,000 SNPs using the *k*-means clustering algorithm. In each cluster, the accession with the highest genotypic value was selected as a parent for a four-way cross. The four-way cross comprised the four selected accessions and yielded 150 individuals from hybridizations between two different F_1_ parents.

### Breeding program

In the breeding program, we adopted the two-part strategy proposed by Gaynor et al. (2017) and adapted it for use in inbred crops that lack established protocols for producing doubled haploids. The program comprises a “population improvement component” and “segregation and fixation component,” as illustrated in Figure 1. The population improvement component aimed to enhance the genotypic value of a population through rapid recurrent genomic selection. The segregation and fixation component involved repeated selfing to segregate and fix alleles. The primary objective of this breeding program was to develop genotypes with high genotypic values in the Inbred8 generation for subsequent release as varieties.

**Figure. 1.**
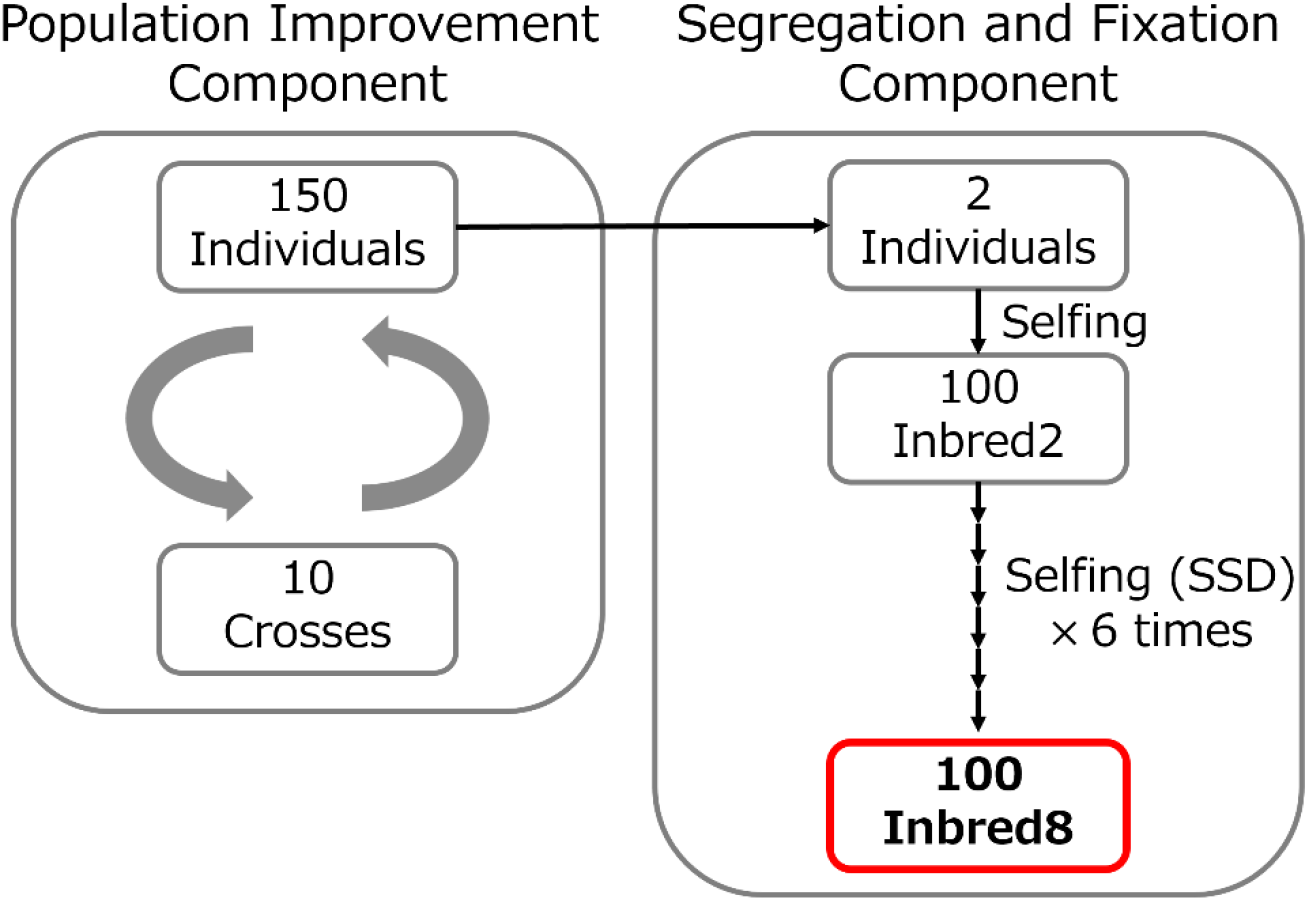
Overview of a breeding program adopted in this breeding simulation. SSD: Single-Seed Decent.

For the population improvement component, each breeding strategy selected 10 crossing pairs from 150 individuals in each cycle (Figure 1). Owing to limitations in the number of flowers and the amount of pollen collected from each soybean plant, we restricted each individual to be used up to twice for crossing pairs in all cycles. Each crossing pair produced 15 progeny, resulting in 150 individuals per cycle. In the subsequent cycle, 10 crossing pairs were selected from a pool of 150 individuals, generating another 150 individuals. This iterative process of selection and crossing enhanced the genotypic potential of a population within the population improvement component.

For the segregation and fixation component, we selected two individuals annually from the 150 individuals in the population improvement component. The selected individuals underwent seven rounds of selfing to segregate and fix the alleles, which fixed >99% of the alleles. The first round of selfing yielded 50 progeny for each individual to facilitate allele segregation, resulting in 100 Inbred2 individuals. Subsequent selfing rounds were conducted using single-seed descent (SSD) method to fix the alleles. Ultimately, 100 Inbred8 individuals were produced annually, and their genotypic values were used for strategy evaluation.

The efficacy of breeding programs depends on the selection and crossing strategies employed in population improvement components. To evaluate the performance of the program, we simulated a 10-year breeding program and compared three breeding strategies (GS, OCS, and CPS). In the population improvement component, selection and crossing cycles were conducted twice a year. Throughout the 10-year breeding program, 20 selection and crossing cycles were carried out, resulting in 100 Inbred8 individuals on 10 occasions. The time required to produce 100 Inbred8 individuals was not considered in this simulation. In practical breeding, the utilization of generation advancement techniques enables a reduction in this timeframe.

### Genomic selection (GS)

This strategy involved selecting 10 individuals with the highest genotypic values, which were calculated using the following formula:

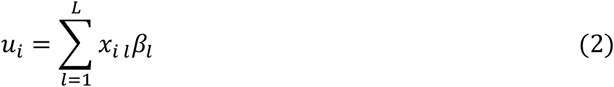

where *u*_*i*_ represents the genotypic value of individual *i*(*i*= 1, …, *N*), *N* is the number of individuals (*N* = 1 5 0), *x*_*i l*_ denotes the SNP marker score of individual *i* on SNP marker *l*, encoded with 0, 1, or 2 for the reference SNP marker, *β*_*l*_ represents the SNP marker effect on SNP marker *l*, and *L* is the total number of SNP markers (*L* = 4, 0 0 0). This strategy selected 10 individuals with the highest genotypic values and randomly determined 10 crossing pairs. Each individual was used twice and duplicate crosses were not permitted.

### Optimal Cross selection (OCS)

This method selects 10 crossing pairs directly from the current population, considering the genotypic values and genetic diversity of the selected individuals (Allier, et al., 2019; Gorjanc et al., 2018). This method involves solving the following optimization problem:

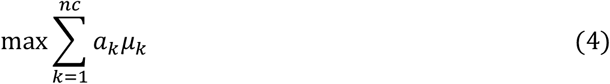

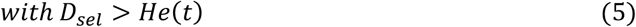

where *nc* is the number of total possible crosses 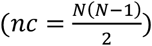, *a*_*k*_ is a dummy variable for each cross (where *a*_*k*_ = 0 indicates that cross *k* is not chosen and *a*_*k*_ = 1 indicates that cross *k* is chosen), *μ*_*k*_ is the mean genotypic value of cross *k*, which can be computed as 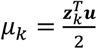, where ***u*** is the genotypic value vector (*N* × 1) calculated in Eq.2, and ***z***_*k*_ is the (*N* × 1) length vector linking the cross *k* to the selected two individuals. *D*_*sel*_ is the genetic diversity of the selected crossing pairs, *He*(*t*) is the genetic diversity constraint at cycles *t* (*t* = 1, …, *T*), and *T* is the final cycle of the population improvement component (*T* = 2 0). *D*_*sel*_ and *He*(*t*) are defined as follows:

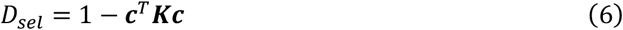

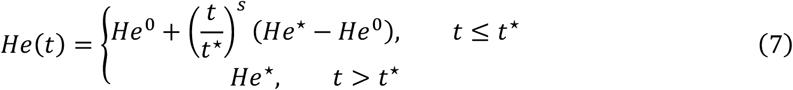

where ***c*** is the individual contribution vector (*N* × 1), computed as 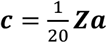 with ***a*** = (*a*_1_, …, *a*_*nc*_)^*T*^ and ***Z*** = (***z***_1_, …, ***z***_*nc*_), ***K*** is the identical-by-state matrix (*N* × *N*), and *He*^0^ is the genetic diversity of the initial population. *t*^⋆^, *s*, and *He*^⋆^ are the parameters in OCS (Allier, et al., 2019). *t*^⋆^ is the target cycle, *s* is the shape parameter, and *He*^⋆^ is the remained genetic diversity in the target cycle. In this breeding simulation, we set *t*^⋆^ = {1 0, 2 0}, *s* = { 0. 5, 1, 2}, and *He*^⋆^ = { 0. 01*He*^0^, 0.1*He*^0^, 0.3*He*^0^}. Eighteen types of OCS models were compared. ***K*** and *He*^0^ were defined as follows:

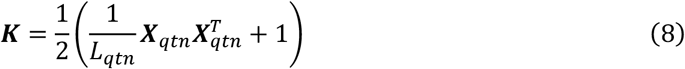

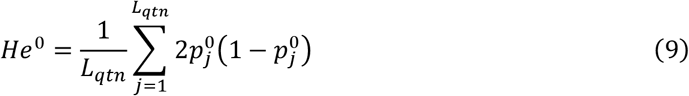

where *L*_*qtn*_ is the number of the QTNs (*L*_*qtn*_ = 1, 0 0 0), ***X***_*qtn*_ is the genotyping matrix (*N* × *L*_*qtn*_), coded in −1, 0, or 1, and 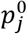 is the frequency of the referent allele in the initial population.

In each cycle, 10 crossing pairs were selected, and each individual could be used up to twice for crossing pairs. The restrictions are defined as follows:

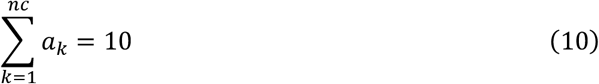

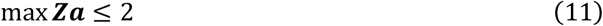

In the OCS, we maximized Eq.4 under Eq.5, 10, and 11. This optimization problem can be solved using the algorithm implemented by Sanchez et al. (2023).

### Cross potential selection (CPS)

The CPS is a novel strategy that selects crossing pairs while considering the segregation of the target generation. The aim of this breeding program was to produce an Inbred8 individual with high genotypic value. Therefore, crossing pairs were selected based on the expected genotypic value of the Inbred8 generation. Following the methodology outlined by Allier et al. (2019), the genetic variance of the Inbred8 population for each cross was computed using the SNP marker effect and score.

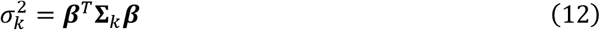

where ***β*** is the SNP marker effect vector (*L* × 1), and **Σ**_*k*_ is the variance-covariance matrix (*L* × *L*), computed from the SNP marker score of cross *k*. Each element of **Σ**_*k*_ represents the variance or covariance between two markers at the Inbred8 generation. Further details on the calculation of this **Σ**_*k*_ matrix for each cross are provided in File S1. In this breeding program, each selected cross produced 15 individuals, and the selected individuals for the population improvement component yielded 50 Inbred8 individuals. When selecting the best individual for the population improvement component from the 15 individuals, the expected maximum value of the Inbred8 individual for each cross was determined as follows:

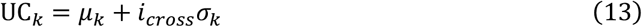

where *μ*_*k*_ is the mean genotypic value of cross *k*, and *i*_*cross*_ ≈ 3. 0 0 4 is a selection intensity that corresponds to selecting the highest Inbred8 individual from the pool of 750 Inbred8 individuals (15 individuals × 50 Inbred8 individuals). Using UC_*k*_, we assessed the potential of each cross to produce outstanding individuals in the Inbred8 population. The objective of the CPS is defined as:

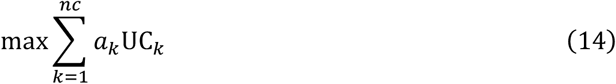

In the CPS, we maximize Eq.14 subject to the constraints outlined in Eq.10 and 11. This optimization problem corresponds to an integer programming problem, which can be addressed using the “lp” function in the “lpSolve” package version 5.6.18 (Berkelaar et al. 2023) in R version 4.1.2.

### Selection for the segregation and fixation component

For the segregation and fixation component, we selected two individuals annually from the 150 individuals in the population improvement component. The same selection method was used in all three strategies. In this breeding program, selected each individual produced 50 Inbred8 individuals. We computed the UC for Inbred8 as follows:

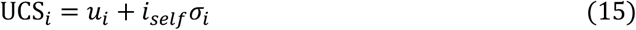

where *u*_*i*_ is the same for Eq.2, *i*_*se lf*_ ≈ 2. 0 5 4 is a selection intensity that corresponds to selecting the highest Inbred8 individual from the pool of 50 Inbred8 individuals, and *σ*_*i*_ is the genetic variance of the Inbred8 population for each individual *i*. Details on the calculation of this *σ*_*i*_ for each individual are provided in File S1. We selected two individuals with the highest UC*S*_*i*_ values for the segregation and fixation component.

### Comparison

We conducted 300 independent breeding simulations for all the three breeding strategies (GS, OCS, and CPS). The primary objective of this breeding program was to generate individuals with high genotypic value in the Inbred8 generation for subsequent release. To compare the effectiveness of the four breeding strategies, we computed the genetic improvements in the Inbred8 population as follows:

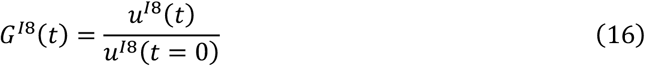

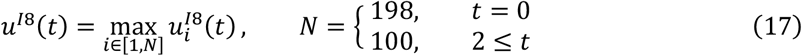

where 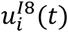 is the genotypic value of individual *i* in the Inbred8 population produced by the two selected individuals from the population improvement component at cycle *t* (*t* = 2, …, 2 0), *N* is the number of Inbred8 individuals, and *u*^*I*8^(*t* = 0) is the maximum genotypic value of a diverse panel of 198 accessions.

The genotypic values of the population improvement component significantly contributed to genetic improvement in the Inbred8 population. Additionally, we computed genetic improvements in the population improvement component as follows:

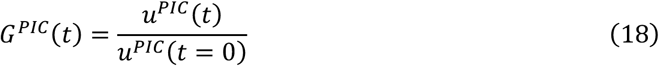

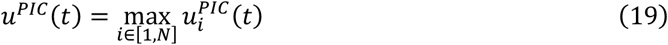

where 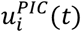 represents the genotypic value of individual *i* in the population improvement component at cycle *t* (*t* = 0, 1, …, 2 0) and *N* is the number of individuals in the population improvement component (*N* = 1 5 0). *u*^*PIC*^(*t* = 0) indicates the maximum genotypic value of the starting population produced from a four-way cross.

Moreover, the genetic variance in the population improvement component serves as a source of genetic improvement. The genetic variance of the population improvement components was computed as follows:

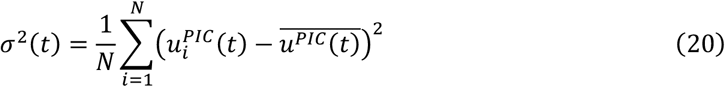

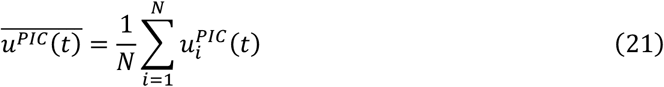

where 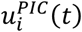 remains the same as in Eq.19.

## RESULTS

### Genetic improvement in the Inbred8 population

Figure 2A illustrates genetic improvements in the Inbred8 generation for each breeding strategy. In OCS, 18 combinations of parameters (*t*^⋆^ = {1 0, 2 0}, *s* = { 0. 5, 1, 2}, and *He*^⋆^ = { 0. 01*He*^0^, 0.1*He*^0^, 0.3*He*^0^}) were tested in 100 independent breeding simulations. OCS1 (*t*^⋆^ = 2 0, *s* = 1, and *He*^⋆^ = 0. 01*He*^0^) showed the best performance in cycle *t* = 1 0, and OCS2 (*t*^⋆^ = 2 0, *s* = 1, and *He*^⋆^ = 0.3*He*^0^) showed the best performance in cycle *t* = 2 0 (Figure S1). Subsequently, the results for OCS1 and OCS2 were used for further analysis. CPS exhibited the highest genetic improvements among the four strategies (GS, OCS1, OCS2, and CPS) in the early cycles (4 ≤ *t* ≤ 1 4) (Figure 2A). Particularly, in cycle *t* = 8, CPS outperformed GS, OCS1, and OCS2 by 6%, 4%, and 5%, respectively. The percentage of CPS showed that the highest genetic improvement at cycle *t* = 8 among 300 independent breeding simulations was 0.73 (Figure 2B). In the late cycles (18 ≤ *t* ≤ 2 0), OCS2 surpassed CPS (Figure 2A), with OCS2 exhibiting a 3% higher genetic improvement than CPS in the final cycle (*t* = 2 0).

**Figure 2.**
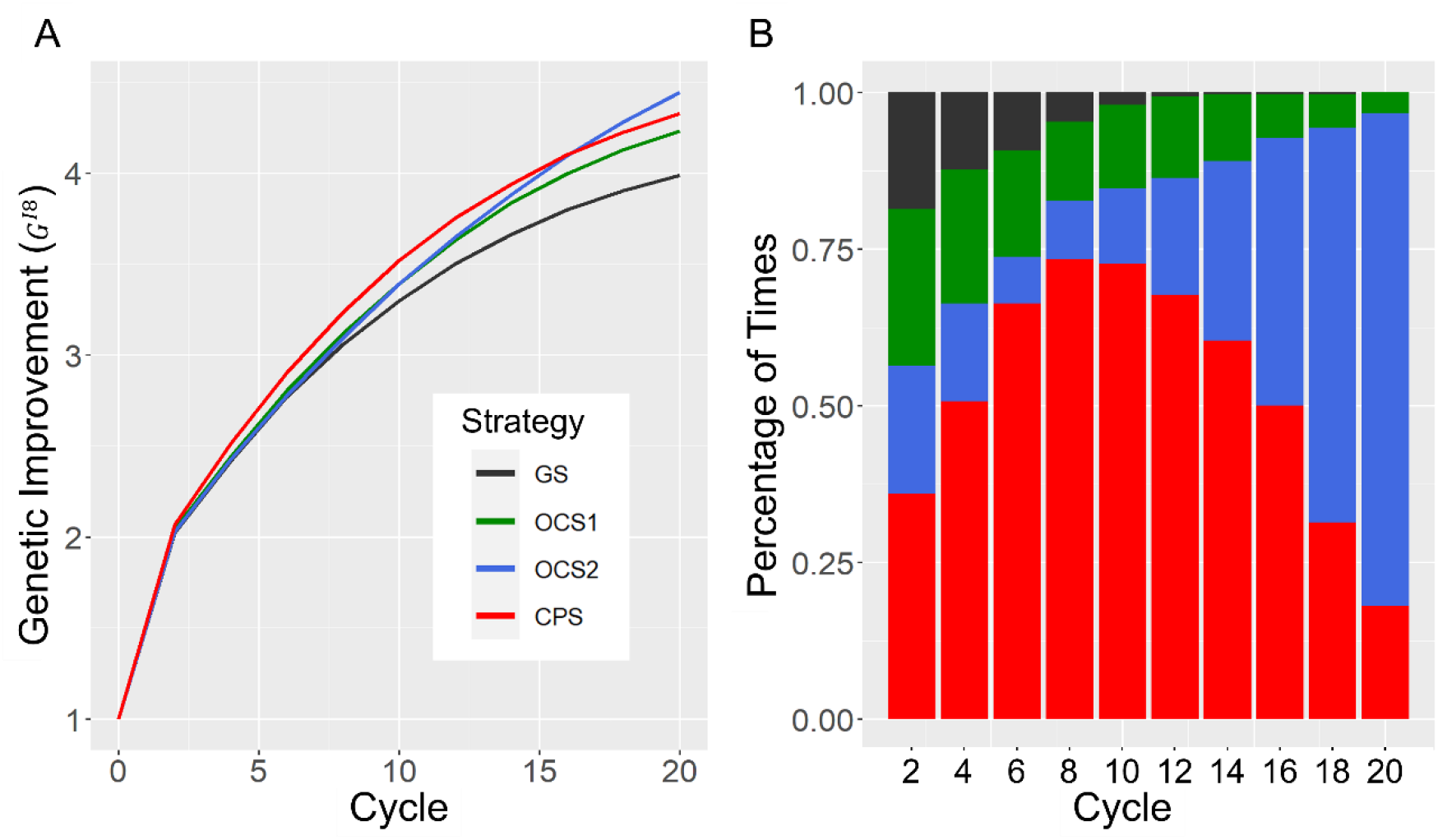
Comparison of four breeding strategies at the Inbred8 generation. GS: genomic selection, OCS1: optimal cross selection (*t*^⋆^ = 2 0, *s* = 1, and *He*^⋆^ = 0. 01*He*^0^), OCS2 (*t*^⋆^ = 2 0, *s* = 1, and *He*^⋆^ = 0.3*He*^0^), and CPS: cross potential selection. (A) genetic improvement (*G*^*I*8^), (B) the percentage of times that each breeding strategy showed the highest genetic improvement for each cycle.

### Genetic improvement and variance in population improvement component

Figure 3A shows the genetic improvement in the population improvement component, which followed a trend similar to that of the Inbred8 generation (Figure 2A). At cycle *t* = 1 0, CPS achieved higher genetic improvements compared to GS, OCS1, and OCS2 by 6%, 4%, and 5%, respectively (Figure 3A). Figure 3B illustrates the genetic variance of the population improvement component. As 150 individuals were produced from one F_1_ cross combination in the initial cycle (*t* = 0) and 150 individuals were produced from 10 crossing pairs from the first cycle (1 ≤ *t* ≤ 2 0), the genetic variance of the first cycle (*t* = 1) exceeded that of the initial cycle (*t* = 0) in all strategies. CPS maintained higher genetic variance than OCS1 and GS over all cycles, whereas OCS2 maintained higher genetic variance than CPS until cycle *t* = 12 (Figure 3B). GS, OCS1, and OCS2 nearly exhausted all genetic variance in the final cycle, whereas CPS retained 27% of the genetic variance from the initial cycle (*t* = 0).

**Figure 3.**
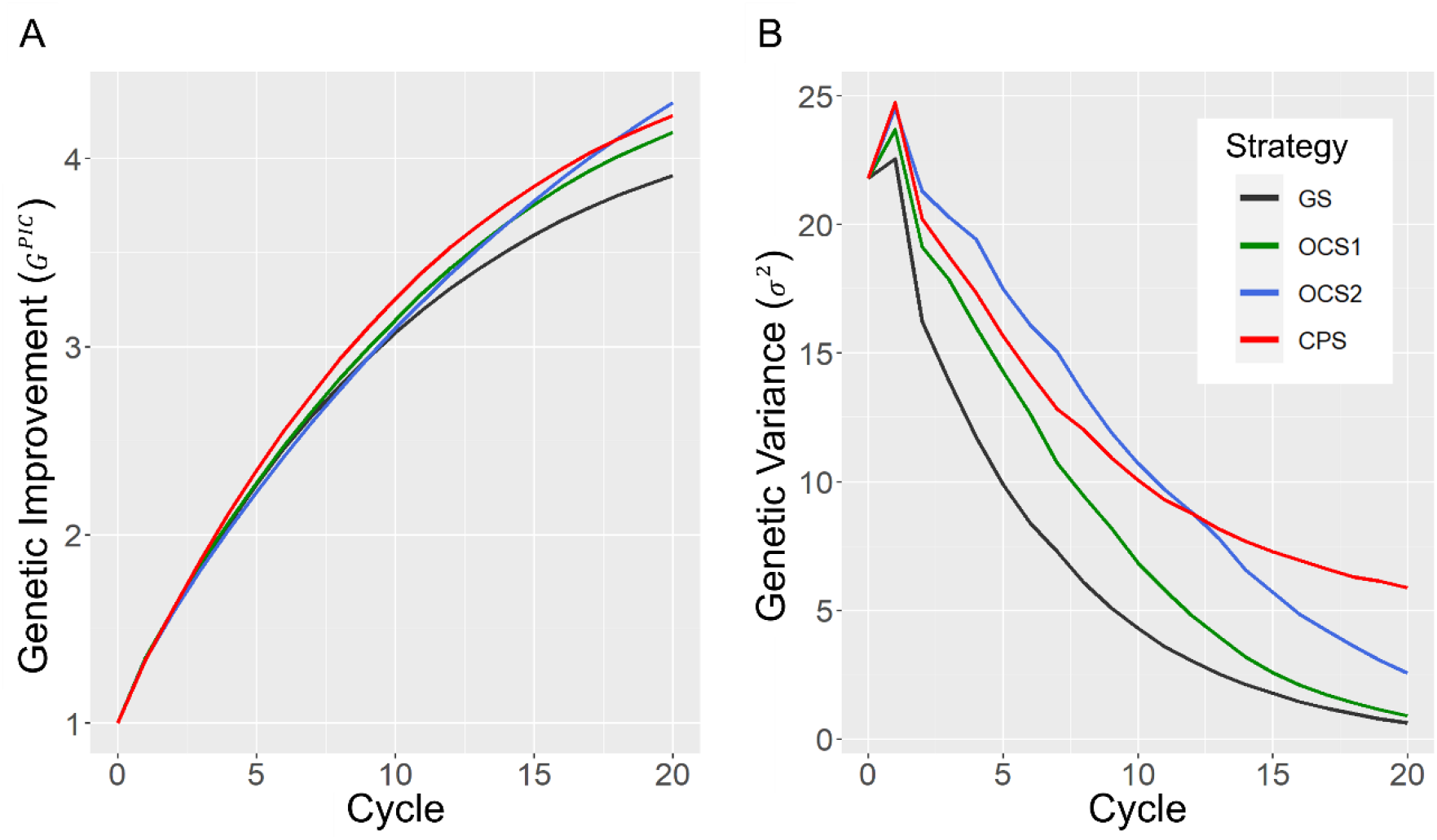
Comparison of four breeding strategies in the genetic improvement component. GS: genomic selection, OCS1: optimal cross selection (*t*^⋆^ = 2 0, *s* = 1, and *He*^⋆^ = 0. 01*He*^0^), OCS2 (*t*^⋆^ = 2 0, *s* = 1, and *He*^⋆^ = 0.3*He*^0^), and CPS: cross potential selection. (A) genetic improvement (*G*^*PIC*^), (B) genetic variance (*σ*^2^).

### Allele states

Figure 4 shows the allele states of the population improvement component for each breeding strategy. The weighted QTN rate of each category (fixed favorable, fixed negative, and non-fixed alleles) was computed as the sum of the absolute effects of the corresponding QTN for each category divided by the sum of the absolute effects of all QTNs. GS fixed numerous alleles compared with the other three strategies (OCS1, OCS2, and CPS) (Figure 4C). OCS2 initially fixed a few alleles and gradually fixed many favorable alleles, targeting the final cycle (Figure 4A, 4C). CPS and OCS1 exhibited a similar trend until cycle *t* = 13, but CPS did not fix as many favorable alleles as OCS1 in the later cycles (1 4 ≤ *t* ≤ 2 0). In the final cycle (*t* = 2 0), the weighted QTN rate of the non-fixed allele in OCS2 was the highest among the four strategies (Figure 4C).

**Figure 4.**
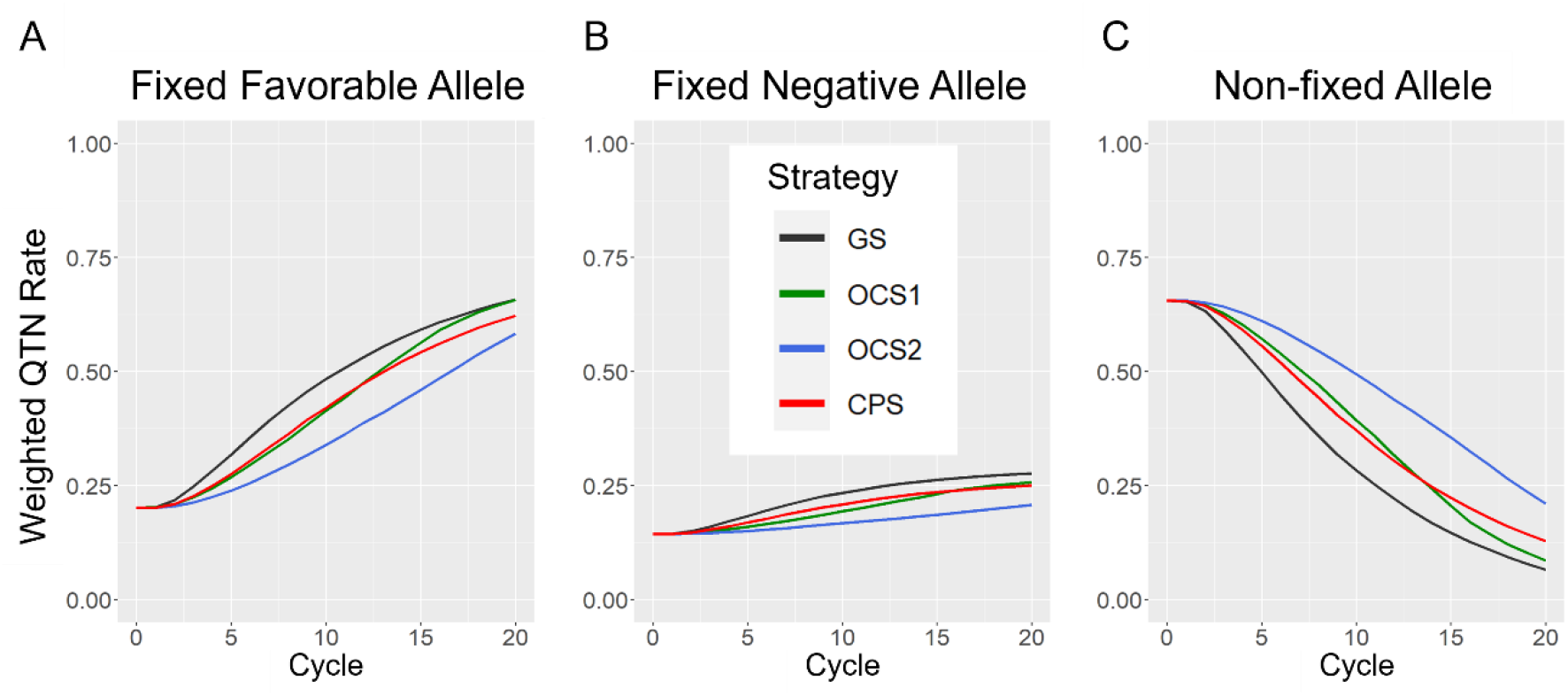
Allele states in the population improvement component for each strategy. GS: genomic selection, OCS1: optimal cross selection (*t*^⋆^ = 2 0, *s* = 1, and *He*^⋆^ = 0. 01*He*^0^), OCS2 (*t*^⋆^ = 2 0, *s* = 1, and *He*^⋆^ = 0.3*He*^0^), and CPS: cross potential selection. The weighted QTN rate of each category is calculated as the sum of the absolute effects of corresponding QTN for each category divided by the sum of the absolute effects of all QTN. (A) fixed favorable allele (B), fixed negative allele (C), non-fixed allele.

## DISCUSSION

### Short-term genetic improvement

In this study, we developed a novel breeding strategy called CPS, which focuses on selecting crossing pairs based on the expected maximum genetic value of the target generation (Inbred8). CPS demonstrated the greatest short-term genetic improvement among the four strategies (GS, OCS1, OCS2, and CPS) (Figure 2). This finding aligns with the outcomes reported by des Déserts et al. (2023), in which crossing pairs selected using UC resulted in superior genetic improvements in the F_5_ population compared to those selected using GEBVs. However, the combination of crossing selection using UC and rapid recurrent genomic selection for rapid varietal development has not been previously explored. Given the imperative to expedite the production of novel varieties through breeding, short-term genetic improvement is essential. In the OCS, adjustments can be made by setting a target cycle (*t*^⋆^) and tuning two parameters (*s* and *He*^⋆^) to achieve high genetic improvement (Allier, et al., 2019; Gorjanc et al., 2018). Despite testing 18 combinations of parameter values (*t*^⋆^ = {1 0, 2 0}, *s* = { 0. 5, 1, 2}, and *He*^⋆^ = { 0. 01*He*^0^, 0.1*He*^0^, 0.3*He*^0^}), no OCS model surpassed the genetic improvement achieved by CPS until cycle *t* = 1 4 (Figure 2, Figure S1). Hence, we posit that CPS is a valuable breeding strategy capable of delivering significant short-term genetic improvements.

### Genetic variance

Genetic variance and diversity in breeding populations are fundamental sources of genetic improvement (Jannink et al., 2010; Sanchez et al., 2023, 2024). OCS effectively maintains genetic variance by integrating a penalty term based on the degree of kinship among selected individuals according to its selection criterion (Allier, Lehermeier, et al., 2019; Gorjanc et al., 2018) In our study, OCS2 consistently maintained high genetic variance and ultimately achieved the most significant genetic improvement among the four strategies by the final cycle (Figure 2, 3B). In contrast, CPS, which did not explicitly consider genetic diversity, managed to sustain a higher genetic variance than OCS1 (Figure 3B). In the CPS, the selection of crossing pairs is predicated on the UC, derived from genetic variance in the Inbred8 generation. As the genetic distance between the two individuals selected as crossing pairs increased, the genetic variance of the Inbred8 population also increased. Consequently, the selection of crossing pairs with greater genetic distances likely contributed to the maintenance of high genetic variance (Figure 3B). The higher heterozygosity observed in CPS than in GS further supports the tendency of CPS to favor crossing pairs with greater genetic distances (Figure 4C). Because of its sustained genetic variance, CPS consistently outperformed GS and OCS1, even in later cycles (Figure 2). Furthermore, CPS retained some potential for additional genetic improvement, as evidenced by its presentation of genetic variance even in the final cycle (*t* = 2 0) (Figure 3B).

However, in an extended 25-year breeding scheme (*t* = 5 0), CPS failed to surpass OCS2 (Figure S2), primarily because of its inability to accurately assess the potential of certain crosses in later cycles. Indeed, in later cycles, the progeny distribution of some crosses deviated from a normal distribution because of the fixation of numerous alleles (Figure S3). Because CPS evaluates each cross under the assumption of a normal distribution of the segregated genotypic values of the progeny, it tends to overestimate the potential of some crosses and selects them as high-potential candidates (Figure S4).

Although CPS may not be optimal for long-term breeding programs, we believe that this overestimation is not a critical issue. Given the focus of our breeding simulations on 1,000 QTNs, it is plausible that real-world scenarios with a larger number of QTNs could provide larger genetic variations for long-term improvement. Additionally, implementing a breeding scheme spanning over 10 years to produce varieties requires significant time and cost. In a 10-year breeding program, CPS accurately evaluated the potential of each cross because the progeny distributions followed normal distributions. Consequently, we believe that CPS is well-suited for short-term genetic improvements, whereas OCS is better suited for long-term genetic improvements.

### Future perspective

In this study, we assessed the efficacy of CPS as a novel breeding strategy for rapid varietal development. Our findings highlight CPS as a valuable approach for achieving short-term genetic improvement, thereby facilitating the expedited production of novel varieties. Although our evaluation employed true marker effects to compare the strategies, it is important to note that in practical breeding programs, the construction of a GP model using training data is essential. Previous research has indicated that the efficiency of strategies such as UC is contingent on the accuracy of progeny variance prediction (des Déserts et al., 2023). Therefore, to effectively implement CPS in real-world breeding programs, it is imperative to develop a highly accurate GP model.

## DATA AVAILABILITY STATEMENT

All datasets and source codes for the breeding simulations are available from the repository in the GitHub, “https://github.com/Sakuraikengo/CPS“.

## ACKNOWLEDGEMENT

The authors thank Dr. Alain Charcosset, Dr. Laurence Moreau, and Dr. Tristan Mary-Huard for their advice regarding this study.

## REFERENCE

Akdemir, D., & Sánchez, J. I. (2016). Efficient breeding by genomic mating. Frontiers in Genetics, 7(NOV), 1–12. 10.3389/fgene.2016.00210

Allier, A., Lehermeier, C., Charcosset, A., Moreau, L., & Teyssèdre, S. (2019). Improving short- and long-term genetic gain by accounting for within-family variance in optimal cross-selection. Frontiers in Genetics, 10(OCT), 1–15. 10.3389/fgene.2019.01006

Allier, A., Moreau, L., Charcosset, A., Teyssèdre, S., & Lehermeier, C. (2019). Usefulness criterion and post-selection parental contributions in multi-parental crosses: Application to polygenic trait introgression. G3: Genes, Genomes, Genetics, 9(5), 1469–1479. 10.1534/g3.119.400129

des Déserts, A. D., Durand, N., Servin, B., Goudemand-Dugué, E., Alliot, J. M., Ruiz, D., Charmet, G., Elsen, J. M., & Bouchet, S. (2023). Comparison of genomic-enabled cross selection criteria for the improvement of inbred line breeding populations. G3: Genes, Genomes, Genetics, 13(11), 1–15. 10.1093/g3journal/jkad195

Diot, J., & Iwata, H. (2023). Bayesian optimisation for breeding schemes. Frontiers in Plant Science, 13(January), 1–14. 10.3389/fpls.2022.1050198

Gaynor, R. C., Gorjanc, G., Bentley, A. R., Ober, E. S., Howell, P., Jackson, R., Mackay, I. J., & Hickey, J. M. (2017). A two-part strategy for using genomic selection to develop inbred lines. Crop Science, 57(5), 2372–2386. 10.2135/cropsci2016.09.0742

Gorjanc, G., Gaynor, R. C., & Hickey, J. M. (2018). Optimal cross selection for long-term genetic gain in two-part programs with rapid recurrent genomic selection. Theoretical and Applied Genetics, 131(9), 1953–1966. 10.1007/s00122-018-3125-3

Jannink, J. L., Lorenz, A. J., & Iwata, H. (2010). Genomic selection in plant breeding: From theory to practice. Briefings in Functional Genomics and Proteomics, 9(2), 166–177. 10.1093/bfgp/elq001

Kaga, A., Shimizu, T., Watanabe, S., Tsubokura, Y., Katayose, Y., Harada, K., Vaughan, D. A., & Tomooka, N. (2011). Evaluation of soybean germplasm conserved in NIAS genebank and development of mini core collections. Breeding Science, 61(5), 566–592. 10.1270/jsbbs.61.566

Kajiya-Kanegae, H., Nagasaki, H., Kaga, A., Hirano, K., Ogiso-Tanaka, E., Matsuoka, M., Ishimori, M., Ishimoto, M., Hashiguchi, M., Tanaka, H., Akashi, R., Isobe, S., & Iwata, H. (2021). Whole-genome sequence diversity and association analysis of 198 soybean accessions in mini-core collections. DNA Research, 28(1), 1–13. 10.1093/dnares/dsaa032

Kinghorn, B. P. (2011). An algorithm for efficient constrained mate selection. Genetics Selection Evolution, 43(1), 1–9. 10.1186/1297-9686-43-4

Lehermeier, C., Teyssèdre, S., & Schön, C. C. (2017). Genetic gain increases by applying the usefulness criterion with improved variance prediction in selection of crosses. Genetics, 207(4), 1651–1661. 10.1534/genetics.117.300403

Meuwissen, T. H. E., Hayes, B. J., & Goddard, M. E. (2001). Prediction of total genetic value using genome-wide dense marker maps. Genetics, 157(4), 1819–1829.

Sanchez, D., Allier, A., Ben Sadoun, S., Mary-Huard, T., Bauland, C., Palaffre, C., Lagardère, B., Madur, D., Combes, V., Melkior, S., Bettinger, L., Murigneux, A., Moreau, L., & Charcosset, A. (2024). Assessing the potential of genetic resource introduction into elite germplasm: a collaborative multiparental population for flint maize. TAG. Theoretical and Applied Genetics, 137(1), 19. 10.1007/s00122-023-04509-5

Sanchez, D., Sadoun, S., Mary-Huard, T., Allier, A., Moreau, L., & Charcosset, A. (2023). Improving the use of plant genetic resources to sustain breeding programs’ efficiency. Proceedings of the National Academy of Sciences, 120(14), e2205780119. 10.1073/pnas.2205780119

Schnell, F. W., & Utz, H. F. (1975). F1-LEISTUNG UND ELTERNWAHL IN DER ZÜCHTUNG VON SELBSTBEFRUfflTERN. In Bericht über die Arbeitstagung der Vereinigung österreichischer Pflanzenzüchter (pp. 243–248).

